# Static stability predicts the continuum of interleg coordination patterns in *Drosophila*

**DOI:** 10.1101/374272

**Authors:** Nicholas S. Szczecinski, Till Bockemühl, Alexander S. Chockley, Ansgar Büschges

**Affiliations:** Department of Animal Physiology, Zoological Institute, University of Cologne, 50674 Cologne, Germany

**Keywords:** motor control, locomotion, insect walking, stability, interleg coordination

## Abstract

During walking, insects must coordinate the movements of their six legs for efficient locomotion. This interleg coordination is speed-dependent; fast walking in insects is associated with tripod coordination patterns, while slow walking is associated with more variable, tetrapod-like patterns. To date, however, there has been no comprehensive explanation as to why these speed-dependent shifts in interleg coordination should occur in insects. Tripod coordination would be sufficient at low walking speeds. The fact that insects use a different interleg coordination pattern at lower speeds suggests that it is more optimal or advantageous at these speeds. Furthermore, previous studies focused on discrete tripod and tetrapod coordination patterns. Experimental data, however, suggest that changes observed in interleg coordination are part of a speed-dependent spectrum. Here, we explore these issues in relation to static stability as an important aspect of interleg coordination in *Drosophila*. We created a model that uses basic experimentally measured parameters in fruit flies to find the interleg phase relationships that maximize stability for a given walking speed. Based on this measure, the model predicted a continuum of interleg coordination patterns spanning the complete range of walking speeds. Furthermore, for low walking speeds the model predicted tetrapod-like patterns to be most stable, while at high walking speeds tripod coordination emerged as most optimal. Finally, we validated the basic assumption of a continuum of interleg coordination patterns in a large set of experimental data from walking fruit flies and compared these data with the model-based predictions.

**Summary statement:** A simple stability-based modelling approach can explain why walking insects use different leg coordination patterns in a speed-dependent way.

## Introduction

Legged locomotion (i.e., walking) is an important behavior for most terrestrial animals; in many species, it is the primary mode of locomotion used in various contexts such as foraging, migrating, finding mates, hunting, or escape. Because of its importance for these behaviors, it can be assumed that walking has become highly optimized during evolution. However, walking is not a fixed behavior and must be adaptable regarding basic parameters like speed or direction. The most prominent of such adaptations is interleg coordination—the temporal and spatial relationship between leg movements. In large vertebrates like dogs, horses, and humans, changes in walking speed are accompanied by changes in interleg coordination, termed *gait* transitions (Alexander, 1989). A gait can be defined as a distinct mode of locomotion used within a particular speed range. For instance, a horse will first walk at low speeds then transition to trot at an intermediate speed and, finally, switch to gallop at high speeds (Orlovsky et al., 1999). The transition between two gaits occurs at a characteristic locomotion speed and is discontinuous regarding at least one parameter associated with walking behavior (Alexander, 1989). It is important to note that gaits are not defined by a particular set of movement parameters but by a discontinuous, rather than gradual, transition.

Interleg coordination during walking has also been studied extensively in arthropods, mainly insects (for reviews see Ayali et al., 2015; Bidaye et al., 2017; Borgmann and Büschges, 2015; Cruse, 1990; Cruse et al., 2009). As in vertebrates, these animals adapt their interleg coordination as they change walking speed (Graham, 1972; Wahl et al., 2015; Wendler, 1964; Wilson, 1966; Wosnitza et al., 2013). Several prototypical patterns have been described in the literature; insects use wave gait coordination at low walking speeds (Hughes, 1952), tetrapod coordination at intermediate speeds, and tripod coordination at high speeds (Strauss and Heisenberg, 1990; Wosnitza et al., 2013). Each of these locomotion modes corresponds to a particular interleg coordination pattern. During wave gait coordination, at most one leg executes a swing phase at any given time, while metachronal waves of protraction progress from the hind to the front leg on each side of the animal’s body. In tetrapod coordination, at most two legs are in swing phase at a particular time. Finally, tripod coordination is characterized by concurrent swing phases of ipsilateral front and hind legs and the contralateral middle leg.

Commonly, these interleg coordination patterns in insects are referred to as gaits in the literature (Bender et al., 2011; Dürr et al., 2018; Nishii, 2000; Ramdya et al., 2017; Spirito and Mushrush, 1979); however, to our knowledge, it has never been explicitly shown that the different forms of locomotion found in insects actually fulfill the definition of gaits as suggested by Alexander (1989)—namely, that these are discrete modes of locomotion and not merely special cases along a continuum.

Based on data from the cockroach *Periplaneta americana* (Hughes, 1952) and the stick insect *Carausius morosus* (Wendler, 1964), Wilson (Wilson, 1966)proposed a set of simple rules for the generation of interleg coordination in six-legged insects. In contrast to the common assumption of actual gaits in insects, these rules predicted that insects should use a speed-dependent continuum of interleg coordination patterns. Wilson also pointed out that these rules should result in the natural emergence of all known coordination patterns, including wave gait-like, tetrapod, and tripod coordination, as part of this continuum. Similarly, Spirito and Mushrush (1979) clearly showed a continuum of phase relationships between legs in walking *P. americana*. Results from *Drosophila melanogaster* support the notion of a continuum of coordination patterns; the tripod coordination strength calculated in a study by Wosnitza et al. (2013) showed no clear discontinuities when analyzed over the complete range of walking speeds.

These studies suggest that walking insects change interleg coordination in a speed-dependent, continuous, and systematic manner and either imply, describe, or explain this continuum. However, to our knowledge there has been no explicit attempt to explain why these changes occur (i.e., what the adaptive value of these changes might be). Tripod coordination, which is typically used at high walking speeds, would also be suitable for slow walking; indeed, fruit flies can also use tripod coordination at lower speeds (Gowda et al., 2018; Wosnitza et al., 2013). However, there is no directly evident reason why insects should shift to different, more tetrapod-like interleg coordination patterns at low speeds. The fact that a tendency for this shift nevertheless can be observed in most insects suggests that some aspect of the shift to other interleg coordination pattern must be more optimal at lower speeds as compared to tripod. Of course, exceptions are known: dung beetles (genus *Pachysoma*), for instance, sometimes use a peculiar galloping gait (Smolka et al., 2013), and *P. americana* can switch to quadrupedal and even bipedal running during high speed escape (Full and Tu, 1990).

In the present study, we explored the question of why walking insects change interleg coordination in a speed-dependent manner. In large animals, energy optimality is typically assumed to be the crucial factor responsible for the emergence of true gaits (e.g., Hoyt and Taylor, 1981). Here, we consider static stability during walking as a potentially important parameter. To investigate the influence of stability on coordination, we devised a compact model that incorporates several kinematic parameters that are known from walking fruit flies (*D. melanogaster*), such as swing duration, stance amplitude, and stance trajectory. Fruit flies spontaneously walk at various speeds, so data from these animals is well suited to explore a large range of walking speeds (Mendes et al., 2013; Strauss and Heisenberg, 1990; Wosnitza et al., 2013). The model was used to exhaustively test all theoretically possible coordination patterns (defined herein as phase relationships between ipsilaterally or contralaterally adjacent legs) for all experimentally observed walking speeds in *Drosophila*. The predicted phase relationships between legs were then compared with a large body of corresponding data from walking flies.

The results herein suggest that static stability plays a role in the selection of interleg phase relationships. At high reference walking speeds, our model predicts that tripod-like coordination is the optimal coordination pattern. This changes when the reference speed is lowered to speeds that, in the fruit fly, are found in the intermediate or slow range; here, legs are less tightly coupled, and the animal takes advantage of more stable coordination patterns. The patterns predicted by the model resemble tetrapod-like and wave gait-like coordination. Importantly, the model predicts a continuum of coordination patterns that smoothly vary with walking speed. Experimental data confirm that walking flies shift their coordination in a similar way; their motor output seems to also reflect not only theoretically attainable stability but also how robustly such stability can be realized in the presence of locomotor variability.

## Materials and Methods

### Stability model

Based on previous experimental findings (Wosnitza et al., 2013) we created a model that incorporates several key aspects of walking in *D. melanogaster* and explicitly addresses the speed-dependent nature of inter-leg coordination. The model makes the following assumptions:

1. The duration of stance phase depends on walking speed.
2. Each leg’s stepping frequency depends on walking speed.
3. The duration of swing phase does *not* depend on walking speed.
4. The stance amplitude does *not* depend on walking speed.
5. The phase relationships between pairs of ipsilateral legs are identical.
6. The phase relationships between pairs of contralateral legs are identical.

Data from a previous study (Wosnitza et al., 2013) validate these assumptions and are presented in Figure 1. Least-squares fitting reveals that swing phase duration and step amplitude are only weakly correlated with walking speed. In contrast, stance phase duration and step frequency are strongly correlated with walking speed. Importantly, both stepping frequency and stance duration can be accurately predicted assuming that swing duration and step amplitude are constant (Fig. 1C and D).

**Figure 1:**
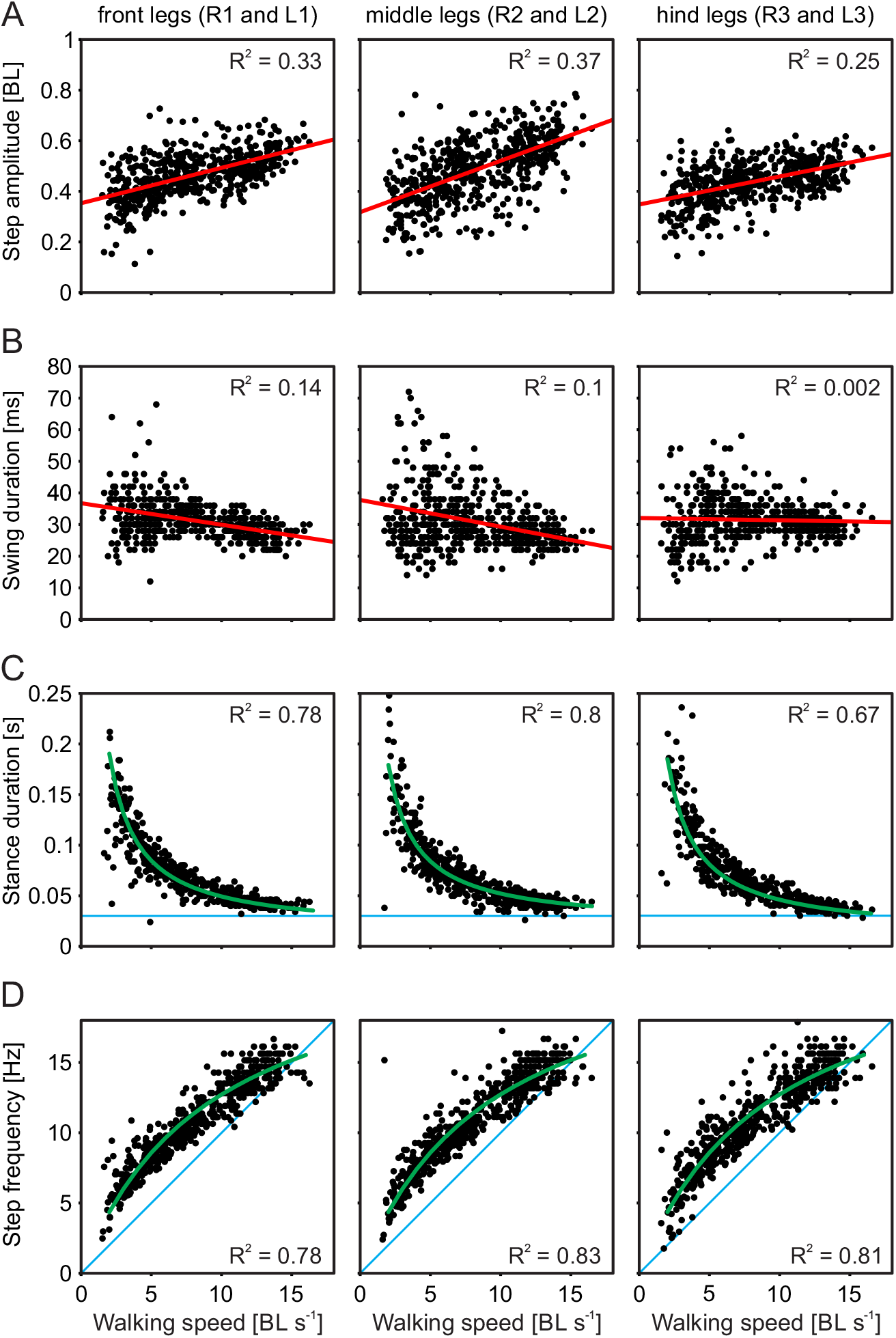
Basic kinematic and temporal parameters of walking *D. melanogaster*. All parameters are expressed as a function of walking speed. Points correspond to individual steps. Left column corresponds to front legs (left and right, L1 and R1), middle column to middle legs (L2 and R2), right column to hind legs (L3 and R3). (A) Step amplitude is only weakly correlated with walking speed (regression line in red). (B) Swing duration is constant over the observed speed range (regression line in red). (C and D) Both stance duration and step frequency are strongly correlated with walking speed. Assuming that step amplitude and swing duration are constant, stance duration and step frequency can be predicted with high accuracy (green lines and corresponding coefficients of determination in C and D). This figure was created with experimental data from Wosnitza et al. (2013).

The model presented here used these relations and a desired walking speed as a set point to calculate the corresponding stepping frequency and stance duration. These two parameters were then used in conjunction with experimentally measured average stance trajectories (Fig. 2C; data from Wosnitza et al., 2013) to construct one complete step cycle for each leg. All stance trajectories were defined in relation to the center of mass (COM) of the fly. The COM’s position was estimated by individual weight measurements of heads, thoraces, abdomina, sets of six legs, and the wings (n = 30). These measurements showed that the head contributed 12.5% of the fly’s total weight, the thorax contributed 31%, and the abdomen 45%. The combined weight of the legs (11%) and the wings (0.5%) were neglected for the calculation of the center of mass. The head, thorax, and abdomen were then modeled as conjoined ellipses that had the same dimensions and relative positions as their counterparts. Using the individual weights and the positions and dimensions of the modeled body parts, we calculated the position of the COM.

**Figure 2:**
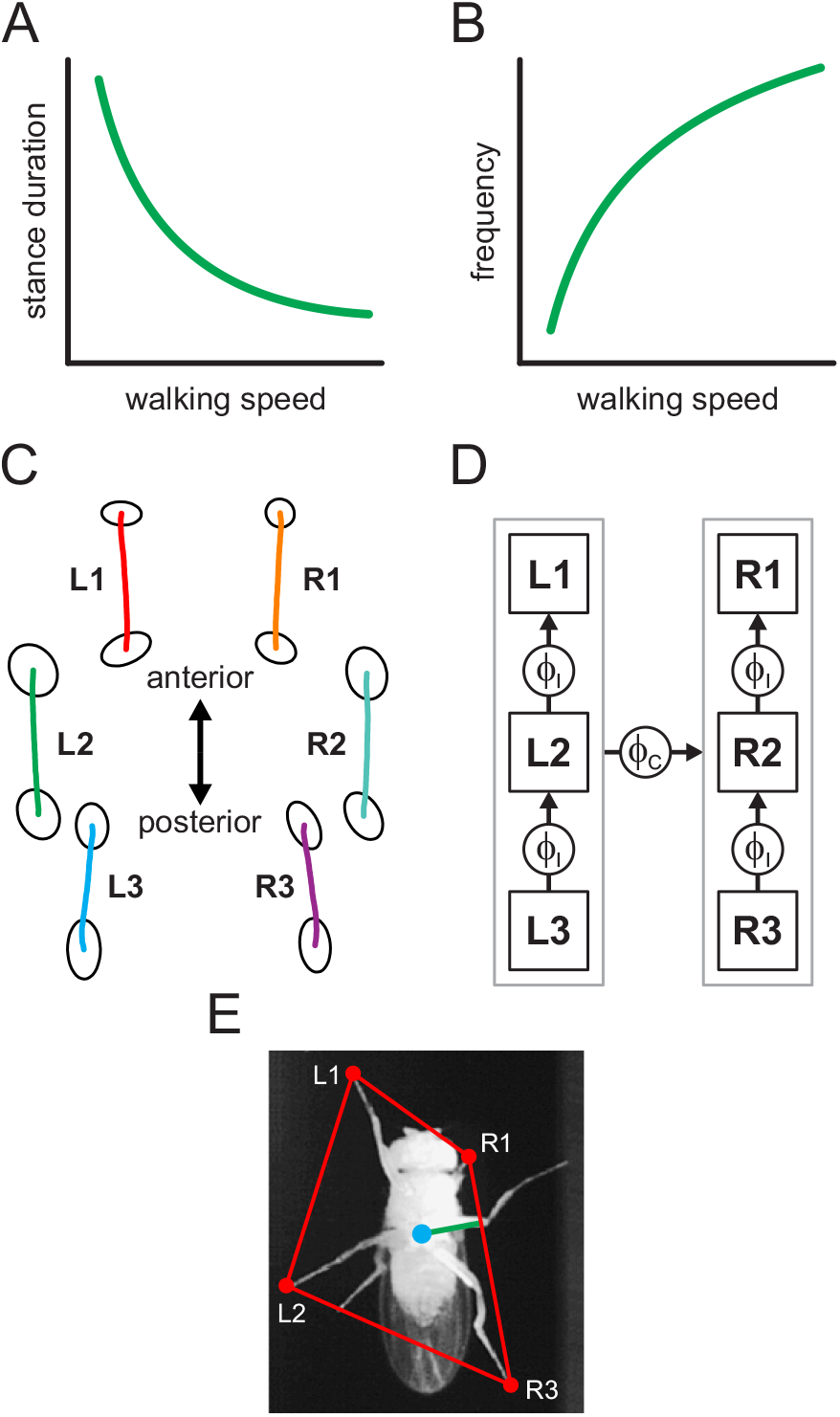
Kinematic model and static stability. (A and B) The predicted relationships between walking speed and stance duration (A) and stepping frequency (B; see Fig. 1C and D), respectively, determines the frequency and stance duration for a particular speed. Thus, a temporal sequence of swing and stance movements can be calculated for each leg. (C) Average stance trajectories measured during experiments are combined with this temporal sequence of swing and stance movements. A stance movement begins at the AEP (anterior-most point in a stance trajectory), progresses with uniform speed (i.e., the set walking speed) to the PEP (posterior-most point in a trajectory), is interrupted by the swing movement, and then starts again from the AEP. Stance trajectories are described in body-centered coordinates. Ellipses around AEPs and PEPs indicate one standard deviation of positional variability measured in experiments (however, only average stance trajectories were used here and variability was not taken into account). (D) Two values, *ϕ_I_* and *ϕ_C_*, describe the phase relationships between ipsilateral legs and contralateral body sides, respectively. Since all other parameters are either constant (stance trajectories) or uniquely defined by a particular walking speed, these two values are the only free parameters in the model. (E) Thus, for a given set of *ϕ_I_* and *ϕ_C_* and at a particular time within one complete step cycle, it can be determined which legs are currently in stance and what their positions with regard to the center of mass (blue dot) are. The legs currently in stance form a convex hull; the minimal distance between the center of mass and the convex hull defines static stability (green line). Its minimum value over one complete step cycle defines the stability for a particular set of *ϕ_I_* and *ϕ_C_*.

During virtual swing movement, a leg’s tarsus was lifted off at the posterior extreme position (PEP) and moved to the anterior extreme position (AEP). During the virtual stance movement, the tarsus touched down at the AEP and moved with a uniform speed (i.e., the set walking speed) to the PEP, where it was lifted off again. Two parameters, *ϕ_I_* and *ϕ_C_*, determined the phase relationships between the legs in this model (Fig. 2D); they can adopt values between 0 and 1. *ϕ_I_* defined ipsilateral phase relationships of step cycles between hind and middle legs and between middle and front legs. Each set of three ipsilateral legs was then treated as a unit (gray outline in Fig. 2D), and the phase relationship between these contralateral units was determined by *ϕ_C_*. Thus, for a particular walking speed and a set of phase relationships, a particular leg’s position and whether it was in stance could be determined at a given time. The tarsal positions of the legs simultaneously in stance at a given time were used to determine a support polygon; the minimum distance between the COM and an edge of this polygon was defined as stability (Fig. 2E). Stability was positive when the COM was within the support polygon and 0 when it was outside. When there were fewer than three legs on the ground, stability was undefined and set to 0.

For a set walking speed, a stepping frequency and stance duration were uniquely defined, and the average stance trajectories were assumed to be constant. Consequently, there were two adjustable parameters in this model: *ϕ_I_* and *ϕ_C_*. To determine stability for different sets of *ϕ_I_* and *ϕ_C_*, each of the two phases was varied systematically from 0 to 1 in steps of 0.02. For each possible combination of phase relationships, we simulated one complete step cycle and calculated its minimum stability. Thus, coordination patterns for which the COM always remained within the support polygon returned positive values. Those with larger values keep the COM towards the center of the polygon at all times, increasing the margin of stability.

### Flies and animal husbandry

Fruit flies (*Drosophila melanogaster*) were raised at a temperature of 25 °C and 65% humidity on a 12-h/12-h light/dark cycle. They were raised on a medium based on a recipe by Backhaus et al. (1984). Experimental data were based on three different fly strains for the experiments described herein: the wild-type strains *Berlin* and *CantonS* (WT, data from this study and Wosnitza et al., 2013) and the mutant strain *w^1118^* (data from Wosnitza et al., 2013). These mutant flies have been reported to walk more slowly than wild-type strains, but show no other apparent impairments (Wosnitza et al., 2013). Flies used during experiments were between three and eight days old. Fly data presented in the manuscript were either obtained during free-walking or tethered walking.

### Free-walking assay

A schematic of the free-walking setup is shown in Figure 3A. It consisted of an inverted glass petri dish that we used as a transparent arena (diameter 80 mm) held by a circular frame with a cutout below the dish. The cutout provided an unobstructed bottom view of the arena.

A surface mirror was placed below the arena at a 45° angle; this allowed for video recordings at approximately the same height as the setup. In conjunction with the mirror, we used an infrared (IR)-sensitive high-speed camera (VC-2MC-M340; Vieworks, Anyang, Republic of Korea) to capture a bottom view of a central rectangular area on the surface of the arena of approximately 30 x 36 mm, with a resolution of 1000 x 1200 pixels, 200-Hz frame rate, and a shutter time of 200 μs. Illumination was provided by a ring of IR light-emitting diodes (LEDs) arranged concentrically around the arena and emitting their light mainly parallel to the arena’s surface. This resulted in a strong contrast between background and fly (see Fig. 3B). The LEDs’ activity was synchronized to frame acquisition of the camera. To prevent escape, the arena was covered with a watch glass that established a dome-shaped enclosure, similar to an inverted FlyBowl (Simon and Dickinson, 2010). To keep flies on the horizontal petri dish, we covered the inside of the watch glass with SigmaCote (Sigma-Aldrich, St. Louis, MO). Prior to an experiment, a single fly was extracted with a suction tube from its vial and placed onto the arena, which was then immediately covered with the watch glass. Flies were allowed to explore the arena for approximately 15 minutes, after which video acquisition was started.

**Figure 3:**
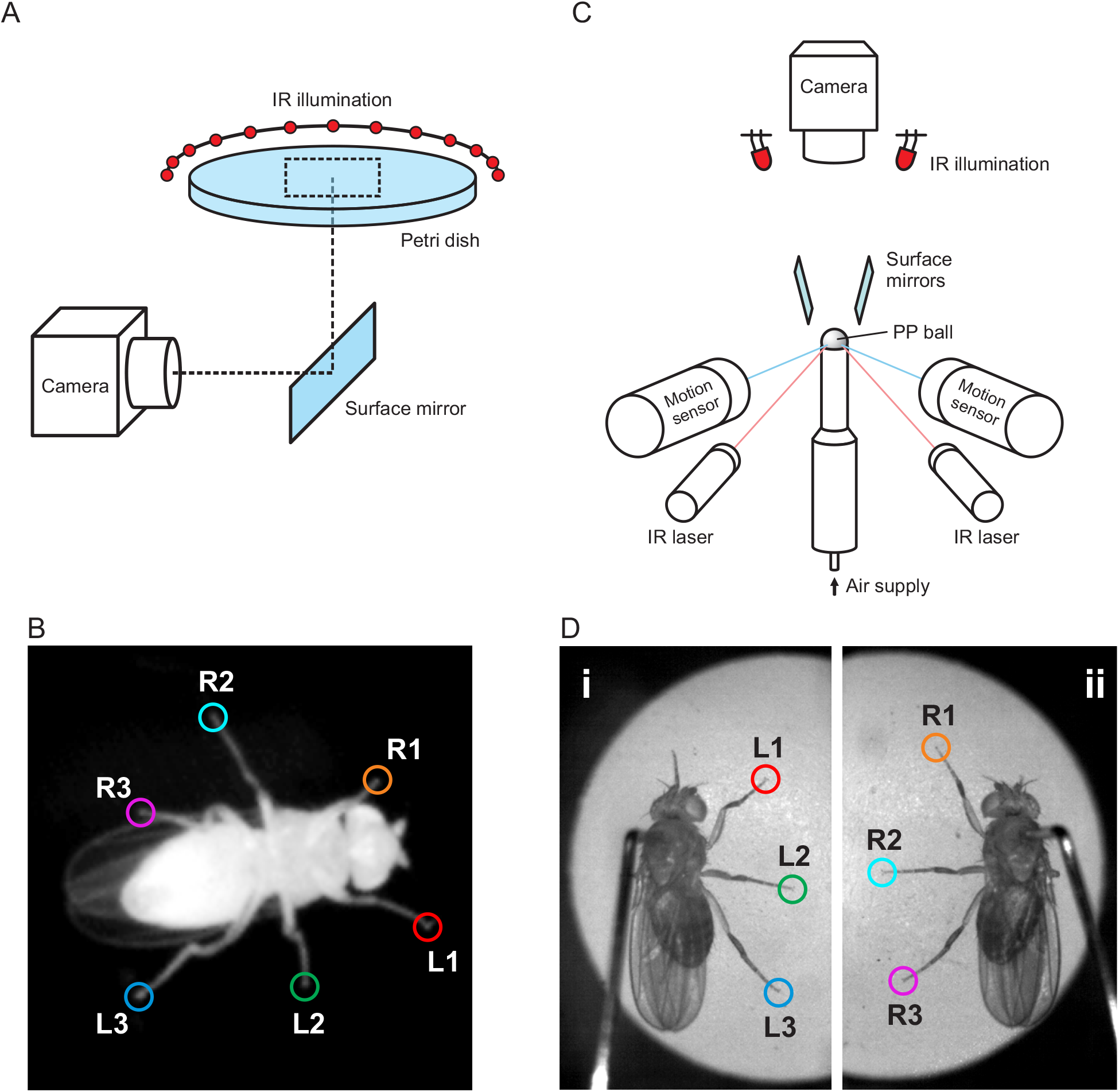
Experimental setups. (A) Free-walking setup. Flies walked on top of a glass petri dish covered with a watch glass (not shown for clarity). A concentric ring of IR LEDs provided illumination (ring only shown partially). A high-speed camera captured a rectangular area of the petri dish (dashed rectangle) via a surface mirror. As soon as the fly walked through the capture area, the recorded video was committed to storage for post-processing. (B) Example cutout from a video frame captured in the free-walking setup. Leg tips are clearly visible and have been manually annotated (for labels see Fig. 1). (C) Tethered-walking setup. Flies walked on an air-supported PP ball whose rotational movements were captured by two motion sensors. Illumination for the sensors was provided by IR lasers. The top of the ball and the two mirrors were captured with a high-speed camera mounted above the setup; illumination for the camera was provided by an LED ring around the camera lens. (D) Two surface mirrors provided side views of the walking fly. Legs are clearly visible in these views; leg tips have been annotated manually.

Flies were spontaneously active in the arena and frequently crossed the capture area. Video data of this area was continuously recorded into a frame buffer of five-to-ten-second durations. During an experiment, custom-written software functions evaluated the recorded frames online and determined if a fly was present and if it had produced a continuous walking track that was at least 10 body lengths (BL) long. Once the fly had produced such a track and either stopped or left the capture area, the contents of the frame buffer were committed to storage as a trial for further evaluation. Video acquisition and online evaluation during acquisition were implemented in MATLAB (2016b; The MathWorks, Natick, MA).

### Tethered-walking assay

A schematic of the tethered walking setup is shown in Figure 3C. It is a modified version of a setup described previously (Berendes et al., 2016; Seelig et al., 2010). The setup consisted of an air-supported polypropylene ball (diameter 6 mm) onto which a tethered fly can be placed. Flies placed atop the ball in this manner will show spontaneous walking behavior and use the ball as an omnidirectional treadmill. Ball movements were measured by two optical sensors (ADNS-9500; Broadcom, Inc., San Jose, CA) with an acquisition speed of 50 Hz. Each of these sensors provided information about 2D optic flow at the equator of the ball; combining these data allowed for the reconstruction of the ball’s rotational movement around its three axes of rotation. Based on these movements, we reconstructed the fly’s instantaneous speed and the curvature of the virtual track during walking. Concurrently, and synchronized to the acquisition of these data, we recorded high-speed video with a resolution of 1200 x 500 pixels from a top view (other parameters and camera model same as above references). Illumination was provided by an IR LED ring positioned around the camera’s lens (96 LEDs) and focused onto the fly. Low-level control of the optical sensors and synchronization to the camera was implemented with custom-made hardware (Electronics Workshop, Zoological Institute, University of Cologne), while high-level control and video data acquisition were implemented in MATLAB. To improve visibility of the fly’s legs, we placed two surface mirrors on a gantry above the fly. The surface of the mirrors formed an angle of 25° with the optical axis of the camera and, thus, provided two additional virtual camera views (see Fig. 3D). Annotation of leg kinematics was done in these side views.

Prior to tethered-walking experiments, flies were cold-anesthetized and transferred into a fly-sized groove in a cooled aluminum block (~4 °C), which held them in place for tethering. Using a dissecting microscope, we then glued a copper wire (diameter 150 μm) to the fly’s thorax. For this, we used dental composite (Sinfony^TM^; 3M ESPE AG, Seefeld, Germany) that was cured within a few seconds with a laser light source (wavelength 470 nm). The wire was inserted into a blade holder which, in turn, was attached to a 3D micromanipulator used for exact positioning of the fly atop the ball. Similar to the free-walking condition, flies were given approximately 15 min to familiarize themselves with the ball and the setup, as well as to recover from anesthesia. Kinematic data from the ball and video data from the camera were captured into separate ring buffers. Flies were spontaneously active; here, however, trial acquisition was done manually.

### Data annotation and analysis

The position of the fly throughout a trial in the free-walking paradigm was determined automatically. In brief, each video frame was converted into a binary image, in which the fly was detected as the largest area. This area was fitted with an ellipse; its major axis and centroid were defined as the fly’s orientation and center, respectively. Walking speed and rotational velocity were calculated as changes of the center and rotation over time. In each trial, the times and positions of all AEPs and PEPs of each leg were determined manually. These positions were then transformed into a body-centered coordinate system based on the fly’s center and orientation. In the tethered-walking assay, walking speed and rotational velocity were provided directly by the ball’s motion sensors. All positional data (speed and distance) were normalized to BL and subsequent analyses were carried out on these body-centered and BL-normalized data.

An individual step was defined as the movement of a leg between two subsequent PEPs. Swing movement was defined as the movement between a PEP and the subsequent AEP; stance movement was defined as the movement between an AEP and the subsequent PEP. The walking speed associated with one step was defined as the average walking speed throughout the step. The instantaneous phase of a step was defined as a value between 0 and 1, which progressed linearly over time between the beginning and the end of the step. The phase relationship between a pair of legs was calculated based on the difference between the instantaneous phases of the two legs at the time of the PEP of one of the legs (i.e., the reference leg). All annotations and calculations were carried out with custom-written functions in MATLAB.

### Results

Our model compactly represents possible interleg coordination patterns (ICP). Figure 4A shows a plot of the ipsilateral phase angle, *ϕ_I_*, against the contralateral phase angle, *ϕ_C_*, which we call a *ϕ*v*ϕ* plot. Each (*ϕ_C_*, *ϕ_I_*) ordered pair represents one ICP. Once a walking speed is set, the full stepping pattern can be determined based on the invariant features we introduced into our model. Figure 4B-E shows several exemplary ICPs corresponding to particular points in the *ϕ*v*ϕ* plot; walking speed was set to 5 BL s^-1^. These examples are meant to give the reader an intuitive understanding of the ¢v¢ plot. For example, when *ϕ_I_* is 1/3, tetrapod-like ICPs emerge (Figure 4B-D). Figure 4B and C illustrate ICPs that have been described in the literature as (ideal) tetrapod patterns. In those, two legs always execute their swing movement at the same time; which legs swing together depends on *ϕ_C_* (either 1/3 or 2/3). As we will show, these ideal tetrapod ICPs are not commonly observed in experimental data, where animals typically use ICPs like the one shown in Figure 4D. The *ϕ*v*ϕ* plot can also describe a tripod ICP (Figure 4E) commonly found in fast-walking insects.

**Figure 4:**
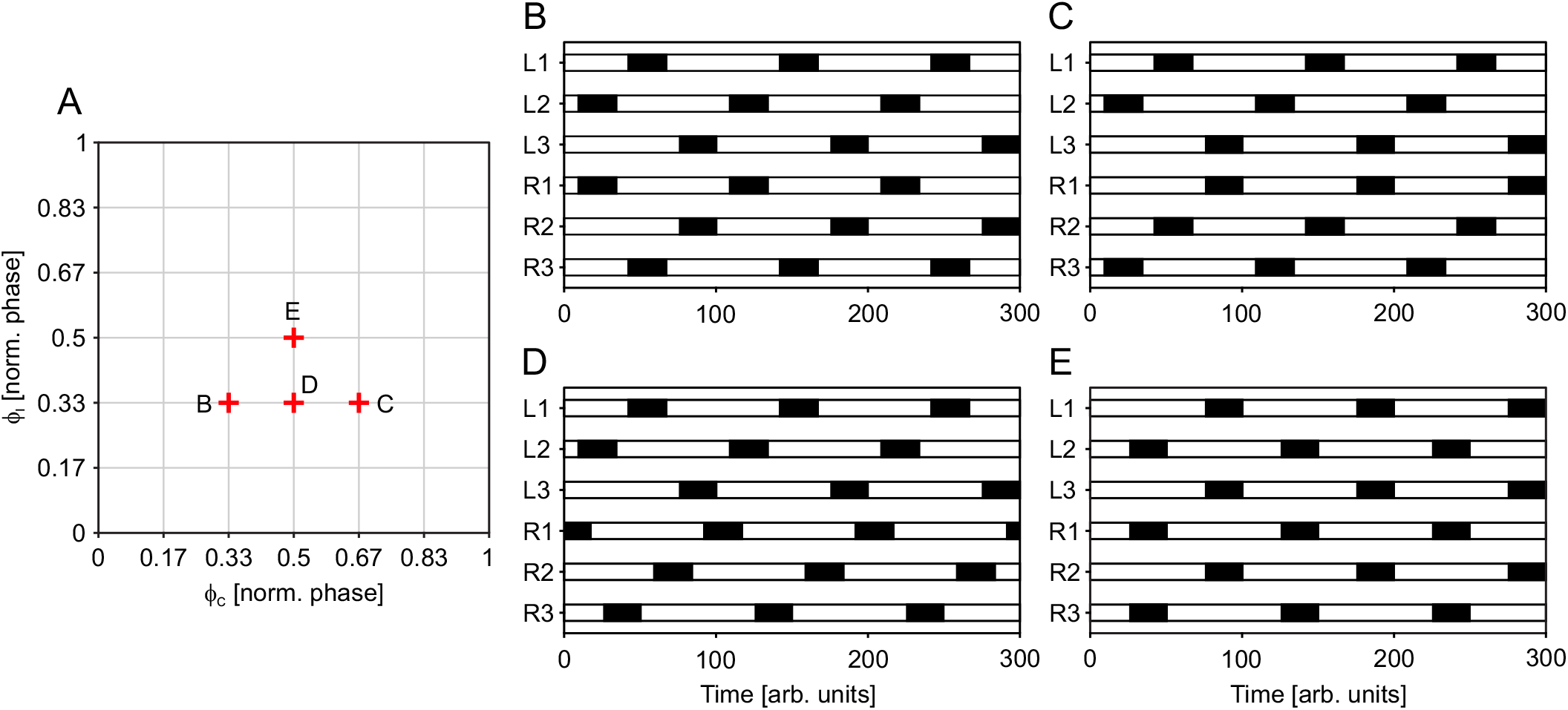
Representative inter-leg coordination patterns (ICPs). Based on the model presented in Fig. 2, each combination of *ϕ_I_* and *ϕ_C_* is associated with a particular ICP. (A) *ϕ*v*ϕ* plot with the position of four exemplary ICPs; each indicated point (B, C, D, E) corresponds to an ICP in panels B to E. (B and C) Idealized tetrapod ICPs commonly referred to in the literature. These correspond to *ϕ_I_* = 1/3 and *ϕ_C_* = 1/3 or 2/3. (D) Tetrapod-like ICP for which *ϕ_I_* = 1/3 and *ϕ_C_* = 1/2. This pattern can be found in walking fruit flies and is also predicted as more stable than the ideal tetrapod ICP (see Results). (E) Tripod ICP corresponding to *ϕ_I_* = 1/2 and *ϕ_C_* = 1/2. This ICP has frequently been reported in the literature and occurs in fast-walking insects. Walking speed for all exemplary ICPs has been set to 5 BL s^-1^ to facilitate comparison. Note that the tripod ICP in E will not occur naturally at these speeds.

The *ϕ*v*ϕ* plots reveal which ICPs are predicted to be the most stable at each walking speed. Figure 5 shows the stabilities of all ICPs at various speeds (Fig. 5Ai-Hi) and the ICPs that correspond to the most stable values of *ϕ_I_* and *ϕ_C_* (Fig. 5 Aii-Hii). Generally, the area showing non-zero stability decreases as walking speed increases. This indicates that, at low walking speeds, more combinations of *ϕ_I_* and *ϕ_C_* result in stable walking. However, unique maxima (i.e., optimal combinations of *ϕ_I_* and *ϕ_C_*) can be found for each walking speed. These phase values of highest stability (red dots in Fig. 5Ai-Hi) indicate that, at low walking speeds, *ϕ_I_* is approximately 0.2 (Fig. 5Ai) and increases continuously towards values of approximately 0.4 (Fig. 5Hi). *Φ_I_* will, in fact, converge to 0.5 at even higher walking speeds (data not shown). At the same time, the optimal value for *ϕ_C_* remains 0.5 over the complete speed range. The footfall patterns associated with the optimal *ϕ_I_* and *ϕ_C_* values in Figure 5Aii-Hii resemble ICPs found in the literature.

**Figure 5:**
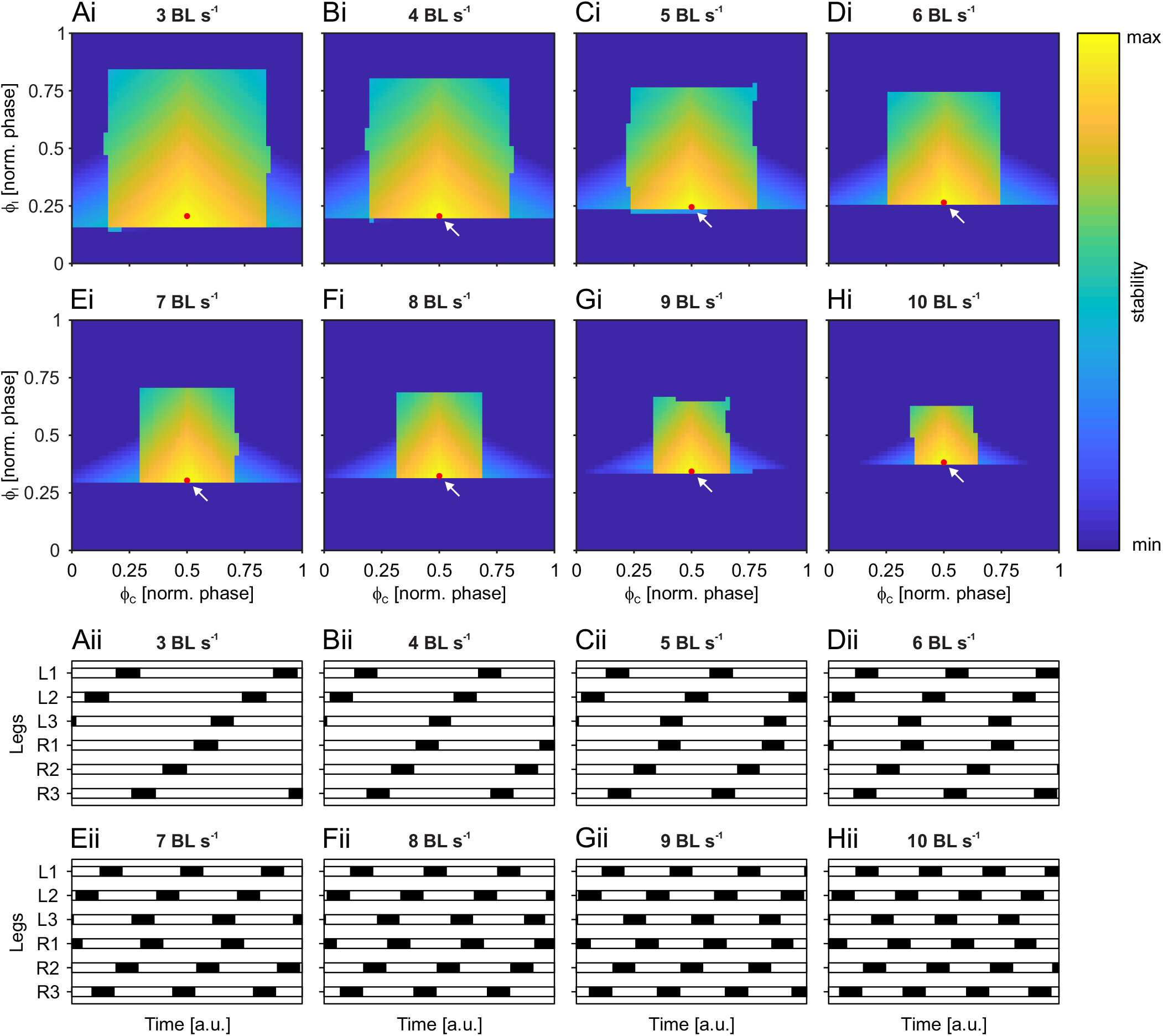
Model-derived stability plots and corresponding ICPs. (Ai to Hi) Each combination of *ϕ_I_* and *ϕ_C_* is associated with a particular stability at a particular walking speed (here 3 to 10 BL s^-1^). High stability is associated with yellow hues, low or zero stability is associated with blue hues. The region of non-zero stability decreases with increasing walking speed and contracts to an area around *ϕ_I_* = 1/2 and *ϕ_C_* = 1/2. In each *ϕ*v*ϕ* plot, the point of maximum stability is indicated by a red dot. Note that at speeds of 4 BL s^-1^ and higher, the points of maximum stability are very close to regions of zero stability (white arrows). (Aii to Hii) ICPs that correspond to *ϕ_I_* and *ϕ_C_* of maximum stability in Ai to Hi. ICPs continuously change from wave gait-like coordination at low speeds to almost tripod coordination at high speeds. Increasing speed to even higher values will, in fact, result in tripod ICPs (not shown).

As walking speed increases, the stance phase duration becomes shorter, reducing the general size of the stable region in each plot. The model predicts that the variance of both *ϕ_I_* and *ϕ_C_* should decrease as walking speed increases, showing an increasingly smaller range of *ϕ_I_* and *ϕ_C_* during the transitions towards tripod. This decrease in variability has been described in the literature and is also apparent in the experimental data presented here (see Fig. 6).

**Figure 6:**
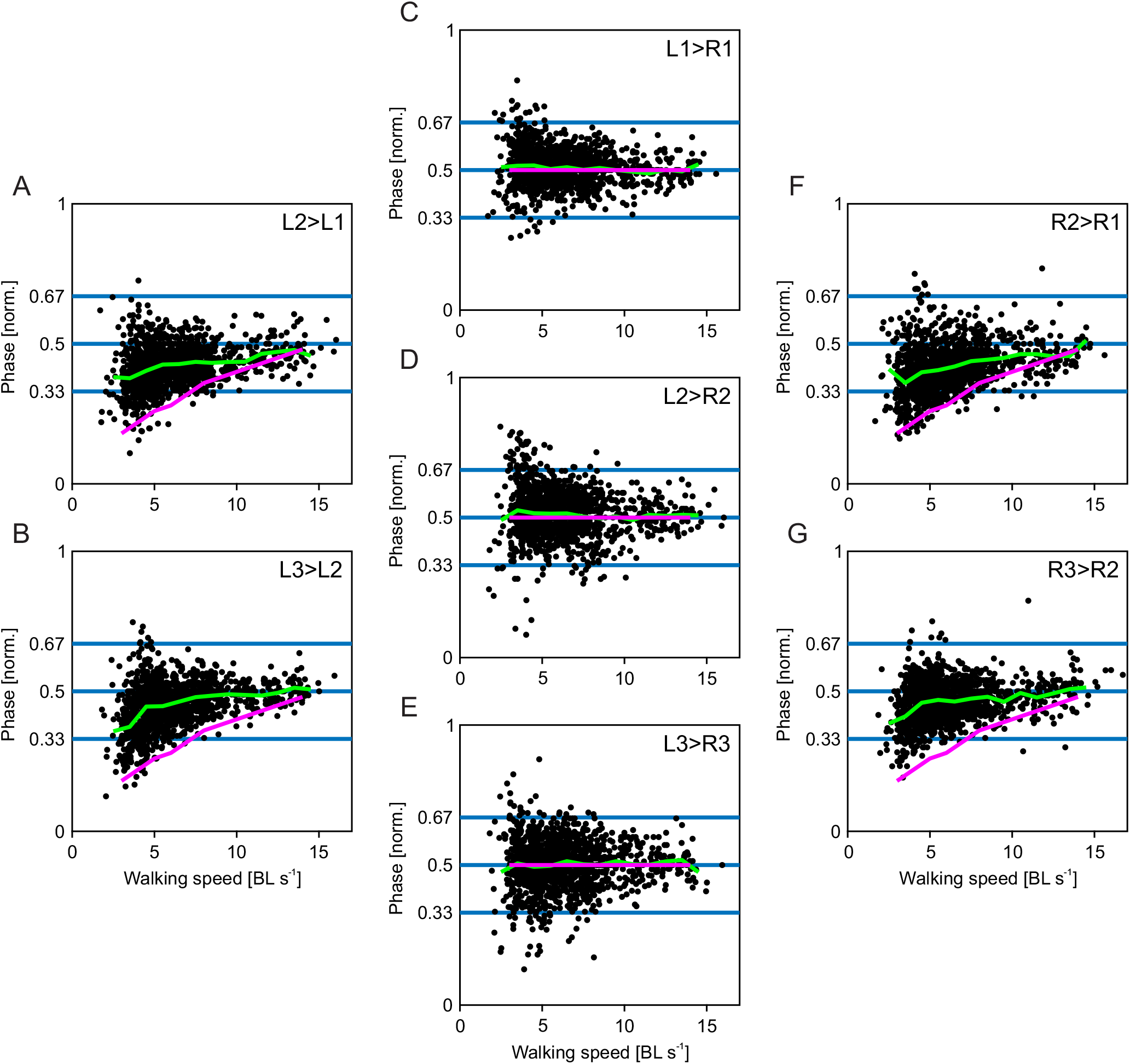
Phase relationships measured during experiments and predicted phases as a function of walking speed. Here, we extended the data set from Wosnitza et al. (2013) with data measured for the present study. Dots correspond to the phase relationship of individual steps, and phase is measured between an observed leg and a reference leg (e.g., L3>L2 refers to the reference leg L3 and the observed leg L2). (A, B, F, G) Phase relationships between ipsilateral middle and front legs (A and F) and hind legs and middle legs (B and G). (C, D, E) Phase relationships between contralateral front legs (C), middle legs (D), and hind legs (E). Green lines indicate running averages of experimentally measured phases; magenta lines indicate model predictions for stability-optimal values of *ϕ_I_* (A, B, F, G) and *ϕ_C_* (C, D, E).

The most stable phase relationships predicted by the model have an anteriorly directed swing phase sequence. This sequence, in which swing phase initiation progresses from the hind leg to the middle leg and ends in the front leg during a complete ipsilateral step cycle, has been described in many studies on six-legged walking in animals, both explicitly and implicitly. The model has not been tuned to adhere to this particular progression; this sequence emerges naturally. Furthermore, as the stability distribution suggests (Fig. 5Ai-Hi), a posteriorly directed sequence, corresponding to *ϕ_I_* values between 0.5 and 1, would be noticeably less stable. This prediction implies a crucial role of the anteriorly directed swing phase progression in walking. It is also noteworthy that the model does not predict the existence of the idealized tetrapod ICP, in which two defined legs simultaneously execute their swing movements. Instead, the model predicts a value of 0.5 for *ϕ_I_* at all walking speeds. The resulting ICPs resemble a tetrapod pattern where, at most, two legs are in swing phase at the same time, but these legs do not enter swing phase simultaneously.

The most stable ICP predicted by the model always lies along the line *ϕ_C_* = 0.5, and its value of *ϕ_I_* depends continuously on the walking speed. To test the model’s predictive ability with regard to these values, we analyzed a pooled dataset (collected in this study and Wosnitza et al., 2013) of 9552 steps (average of 1592 per leg). For 4372 contralateral comparisons and 5849 ipsilateral comparisons *ϕ_I_* and *ϕ_I_* were well defined; in total, we analyzed 106 trials in 31 individuals. We limited our comparison with the model to steps that were produced at walking speeds between 3 and 10 BL s^-1^ during straight walking.

Figure 6 compares stability-optimal values from the model (red lines) with experimental data. Average contralateral phase relationships cluster around 0.5 (green lines, Fig. 6C-D) over the whole speed range, while average ipsilateral phase relationships increase smoothly from values of approximately 0.35 to 0.5 (green lines, Fig. 6A-B, 6F-G). The predicted contralateral phases are very similar to average experimental data (Fig. 6C-D, red and green lines). In addition, the experimental data’s variability decreases towards higher walking speed, which might reflect the reduction in the range of values with non-zero stability (see Fig. 5Ai to Hi). The predicted ipsilateral phases differ noticeably from average experimental data; predicted phase values for *ϕ_I_* are lower than the experimental data. There is, however, a clear tendency towards lower phase values at lower walking speeds. Interestingly, the experimental data seem to be constrained by the optimal phase values predicted by the model at lower speeds, with almost no values below this lower boundary. Figure 5 reveals that the most stable *ϕ_I_* are very close to values associated with low stability or even instability (white arrows). Intuitively, these values correspond to swing movement overlap in ipsilateral neighboring legs (i.e. between hind and middle, or middle and front leg, respectively); any perturbation in the ipsilateral phase relationship that shifts *ϕ_I_* to this lower value will therefore drastically reduce stability. As a consequence, the most stable ipsilateral phase is also the least robust; a small reduction in the ipsilateral phase would destabilize the animal’s posture noticeably. Therefore, the animal appears to prefer more robust ICPs to the most stable ICP. This, in turn, is also evident in the contralateral phase angle data, in which the most stable ICP is also the most robust and the animal can realize this exactly.

One should also note that the model does not predict the existence of the idealized tetrapod ICP, in which two predetermined legs simultaneously execute their swing movement. Instead, the model predicts a value of 0.5 for *ϕ_C_* at all walking speeds. The resulting ICPs resemble a tetrapod pattern (i.e. at most two legs are in swing phase), but these legs do not enter swing phase simultaneously. Discrete changes in gait, like those observed in walking vertebrates, would be apparent as discontinuities in the experimental phase relationships; none are obvious, though, indicating continuous transitions between ICPs.

### Discussion

A large body of data shows that walking at high speeds is associated with tripod coordination in insects, while tetrapod-like and wave gait-like coordination patterns are more frequent at lower speeds. The present work questions why insects change their interleg coordination during walking in such a speed-dependent manner. To address this, we created a stability-based model (Fig. 2) for predicting ICPs during walking in six-legged insects. The model takes into account basic kinematic parameters (Fig. 1 and Fig. 2C) found in walking fruit flies and explicitly accommodates walking speed as an important aspect. Using this model, we exhaustively explored ipsilateral and contralateral interleg phase relationships over the complete range of walking speeds and analyzed the influence of these phases on static stability (Fig. 5). Furthermore, we compared the predicted optimal phase relationships to a large body of experimental data measured in the present as well as a previous study (Wosnitza et al., 2013). The results suggest that stability plays an important role in the selection of an ICP at a particular speed. The model predicts several experimentally observed aspects of insect walking. First, ICPs form a continuum spanning the complete range of walking speeds. Furthermore, it predicts constant contralateral phase relationships of 0.5 and a speed-dependence of ipsilateral phase relationships. The model also provides a potential explanation for the experimentally observed reduction in phase variability at high walking speeds, namely the reduced range of phase values that provide non-zero stability. Finally, an anteriorly directed progression of swing phases in ipsilateral legs emerges in the model.

### ICPs change continuously with walking speed

The model predicts an animal’s preferred ICP at each speed, albeit with some systematic deviation. Furthermore, the speed-invariant contralateral phase angle is predicted to be 0.5, which is also observed in experimental data. The model’s prediction of the ipsilateral phase angle represents one boundary in the experimental data and a sharp edge of stability for the model. This suggests that the animal does not use the most stable ICP, but instead prefers slightly less stable but more robust ICP at a given speed. Regardless, the animal does prefer ICPs that are more stable than tripod, demonstrating that the thoracic ganglia do not function as a centralized tripod generator. Instead, it is likely that a combination of central neural mechanisms and mechanical influences contribute to the animal’s variable, adaptive locomotion.

Our model predicts continuous transitions between ICPs as the walking speed changes, suggesting that fruit flies, and by extension other insects, may not exhibit true gaits like those observed in vertebrates; gait transitions would manifest as discontinuities in such a speed-dependent analysis. Indeed, the experimental data that we collected also showed no evidence of discontinuities indicative of gait transitions. We believe that these data, and those from previous studies in *Drosophila* (Berendes et al., 2016; Wosnitza et al., 2013), support abandoning the term *gait* when referring to insect ICPs, because insect interleg coordination does not fall into discrete coordination patterns. Instead, insect ICPs may be thought of as a continuum of stance durations (Dürr et al., 2018). Based on these findings, we would like to emphasize that walking speed has a strong influence on the parameters measured here (phase relationships and footfall patterns). Studies investigating walking in insects should, therefore, explicitly take into account and measure walking speed to avoid conflating true changes in walking parameters between experimental conditions with mere changes in walking speed.

### Idealized tetrapod ICPs are not preferred

Both our model and the data we collected suggest that *D. melanogaster* does not utilize the idealized tetrapod ICP, in which three pairs of legs sequentially enter swing phase together. While our model suggests that the idealized tetrapod with (*ϕ_C_*, *ϕ_I_*) = (1/3, 1/3) should be a stable ICP (see Fig. 4B and C, as well as Fig. 5), it would be less robust than the observed ICP where (*ϕ_C_*, *ϕ_I_*) = (1/2, 1/3). This is because small changes to either *ϕ_I_* or *ϕ_C_* from (*ϕ_C_*, *ϕ_I_*) = (1/3, 1/3) would destabilize the animal, whereas *ϕ_C_* would have to change substantially from (*ϕ_C_*, *ϕ_I_*) = (1/2, 1/3) to destabilize the animal. Previous studies of walking in *D. melanogaster* have also reported that contralateral legs remain in antiphase at all walking speeds, never giving rise to the idealized tetrapod gait (Strauss and Heisenberg, 1990). Keeping contralateral legs in antiphase at all speeds is also consistent with behavioral descriptions of arthropod interleg coordination (Cruse, 1990) and could potentially simplify interleg control.

Insect interleg coordination is likely determined by more than just the static stability over the course of one step cycle, because the model predicted more extreme speed-dependent changes in ICP (Fig. 5). This discrepancy might be explained by considering the robustness of the coordination pattern—that is, how much error in the interleg phasing can be tolerated before destabilizing the body. By this measure, our model would predict that the animal uses tripod coordination at all speeds, because the stability surfaces in Figure 5 are centered on (*ϕ_C_*, *ϕ_I_*) = (1/2, 1/2). The data suggest that the animal instead utilizes a compromise between the most stable and most robust ICP at a given walking speed, showing variation in the ICP but avoiding potentially unstable ICPs. In fact, the mean (*ϕ_I_*, *ϕ_C_*) of the animal data always lies near the 80^th^ percentile of stable ICPs (data not shown). This means that 20% of other available ICPs would be more stable. However, the animal uses them less frequently, presumably because they are closer to unstable phase relationships. In our comparison between model and experimental data (Fig. 6), the predicted most stable ipsilateral phases (red line) seem to constitute a lower bound for the experimental data; this observation is compatible with the hypothesis that the motor output reflects the expected variability.

### Extensions of the model

Although our model successfully captured the experimental data collected for this study, there are different locomotion scenarios that could be used to test this model in the future. These fall into two main categories: support polygon variant and gravity vector variant. Support polygon variant scenarios include animals with amputated legs and curved walking. In this study, we restricted analysis to intact animals, walking with a very low curvature. However, removing legs drastically affects the support polygon and leads to noticeable changes in ICP in both fruit flies (Wosnitza et al., 2013) and cockroaches (Delcomyn, 1971; Hughes, 1957). In addition, the stance trajectories of fruit fly walking along a curved path are markedly different than during straight walking (Szczecinski et al., 2018). This also changes the associated support polygon and, as a consequence, stability.

Gravity vector variant scenarios include animals walking on inclined, vertical, or inverted substrates. In such cases, the animal is not trying to prevent falling directly toward the substrate as in level locomotion, but at some angle to it, along it, or away from it, respectively. Maintaining stability in such cases would benefit from or require adhesive forces between the animal’s foot and the substrate. In fact, larger insects, such as stick insects, appear to use such mechanisms to improve stability even when walking on flat substrates (Gorb, 1998; Paskarbeit et al., 2016). Studies of insect-inspired climbing robots have shown that the stability of climbing can be analyzed in a very similar way to how we analyzed the stability of walking here, but with the addition of a force tangential to the substrate, provided by the “uphill” leg (Daltorio et al., 2009). In the future, we will expand our model and test its ability to predict ICPs of climbing fruit flies.

### Possible mechanisms in the animal

The goal of this work was not to explain how the animal generates different ICPs, but why. However, it is worth considering which mechanisms may give rise to the phenomena measured herein. Behavioral rules that describe interleg coordination in arthropods have long been known (Cruse, 1990; Dallmann et al., 2017; Dürr et al., 2004). Several of these behavioral rules explicitly address the temporal coordination between onsets of the swing phases in adjacent legs (Rules 1-3, see Dürr et al., 2004). As a consequence, they ensure that the probability of two adjacent legs executing their swing movements simultaneously is low. Recent work with stick insects has shown that the onset of swing phase in a middle leg correlates very tightly with the onset of stance phase in the ipsilateral hind leg (Dallmann et al., 2017). The authors suggest that this is due to the middle leg measuring a decrease in the load being supported, causing the leg to enter swing phase. Indeed, campaniform sensilla, which sense cuticular strain induced by load changes, have been found to be sensitive to unloading in the cockroach (Zill et al., 2009). Such a mechanism could be seen as an indirect measurement of the animal’s stability affecting their ICP. Whether this plays a role in *D. melanogaster*, a particularly light animal, in which gravitational forces might not play a very large role, remains to be investigated.

There is also evidence that walking in insects is more determined by centrally generated motor output at high walking speeds while the influence of leg sensory structures is reduced (Bender et al., 2011; Cruse et al., 2007). This is further supported by recent studies on *C. morosus* (Mantziaris et al., 2017), *C. gregaria* (Knebel et al., 2017), and *D. melanogaster* (Berendes et al., 2016). These studies have shown that neighboring legs have preferred phases of oscillation, even when local sensory feedback is absent. This reduced sensory influence at high walking speeds could, in turn, make the motor output less variable, thus facilitating the convergence to the narrow range of stable ICPs. Ultimately, interleg coordination likely arises through a combination of mechanisms that are mediated both mechanically and neurally.

Assuming that static stability plays a role for the speed-dependent selection of an ICP, an important question is whether stability, or some related proxy, is measured acutely and continuously during walking or if the evolutionary pressure to remain upright that resulted in interleg coordination rules that keep the body upright. Our experimental data from tethered animals whose bodies were supported during walking did not noticeably vary from those from freely walking individuals. In principle, these animals cannot fall over and acute measurement of stability would result in different ICPs. These observations suggest that walking flies do not measure stability as directly as mammals do, for example, utilizing vestibular input (Buschmann et al., 2015).

The consequences of falling are less severe for a fruit fly than for larger animals (Hooper, 2012); if they do misstep and fail to support their body, their large damping to mass ratio should slow down their fall more than for larger animals, such as humans. Nevertheless, fruit flies still need to stay upright during walking. Falling impedes the animal’s progress and wastes energy and time, suggesting that it would benefit the animal to remain upright. This might be even more critical during behaviors like courtship, during which males chase females in close pursuit (Hall, 1994); falling over in this situation might reduce the chances of mating. A similar line of argument can be made for escape from predators, in which precise and smooth stepping is required (Parigi et al., 2014). Stability and the need to remain upright have likely influenced the evolution of the observed ICPs in insects.

## List of symbols and abbreviations

*ϕ_I_*: Ipsilateral phase relationship
*ϕ_I_*: Contralateral phase relationship
AEP: Anterior extreme position
BL: Body length
COM: Center of mass
ICP: Interleg coordination pattern
IR: Infrared
LED: Light-emitting diode
PEP: Posterior extreme position
PP: Polypropylene
*w^1118^*: *D. melanogaster white* mutant strain
WT: *D. melanogaster* wildtype strains *Berlin* and *CantonS*

## Acknowledgements

The authors would like to thank Michael Dübbert and Jan Sydow for excellent technical support, Corinna Rosch for animal husbandry, and Sima Seyed-Nejadi, Sherylane Seeliger, and Hans-Peter Bollhagen for lab support.

## Author contributions

N.S.S., T.B., and A.B. conceived the study, N.S.S., T.B., and A.S.C. carried out the experiments, N.S.S. and T.B. created the model, N.S.S., T.B., and A.S.C. analyzed experimental data, N.S.S., T.B., A.S.C., and A.B. wrote the manuscript. All authors read and approved the final manuscript.

